# Global identification of functional microRNA::mRNA interactions in *Drosophila*

**DOI:** 10.1101/395335

**Authors:** Hans-Hermann Wessels, Svetlana Lebedeva, Antje Hirsekorn, Neelanjan Mukherjee, Uwe Ohler

## Abstract

MicroRNAs (miRNAs) are key mediators of post-transcriptional gene expression silencing. Although *Drosophila* has been of critical importance for miRNA discovery, biogenesis and function, there has been no comprehensive experimental annotation of functional miRNA target sites. To close this gap, we generated the first *in vivo* map of miRNA::mRNA interactions in *Drosophila melanogaster*, making use of crosslinked nucleotides in Argonaute (AGO) crosslinking and immunoprecipitation (CLIP) experiments that enable an unambiguous assignment of miRNAs to AGO binding sites at much higher signal-to-noise ratio than computational predictions alone.

Absolute quantification of cellular miRNA levels showed the miRNA pool in *Drosophila* cell lines to be more diverse than previously reported. Benchmarking two different CLIP approaches, we identified a similar predictive potential to unambiguously assign thousands of miRNA::mRNA pairs from AGO1 interaction data at unprecedented depth. Quantitative RNA-Seq and subcodon-resolution ribosomal footprinting data upon AGO1 depletion enabled the determination of miRNA-mediated effects on target expression and translation. We thus provide the first comprehensive resource of miRNA target sites as well as their quantitative functional impact in *Drosophila*.

## Introduction

MiRNAs are a class of ∼22 nucleotide (nt) long small non-coding regulatory RNAs, involved in mRNA destabilization and translational control. In most cases, a miRNA functions as a guide directing AGO proteins via RNA-RNA-recognition to complementary target sites in the 3’ untranslated region of its target mRNA, where its repressive function gets exerted via assembly of the RNA-induced silencing complex (RISC) ^1^. As miRNAs are predicted to target more than 50 % of all 3’UTRs of protein coding genes in human ^2^ and 30% of Drosophila genes (TargetScanFly 6.2), either as a single miRNA or in combination, they may be the most prevalent negative regulator of post-transcriptional gene expression.

Historically, *Drosophila melanogaster* has been an important model organism to study miRNAs biogenesis and function ^3,4^. MiRNA gene *null* flies identified miRNAs that are critical for fly development as negative regulators of the anti-apoptosis genes *hid* (bantam) and *Drice* (miR-14) ^5,6^. Many fly miRNAs exhibit spatial and temporal expression patterns and possibly regulation ^7–9^. Advances in detecting miRNAs and their systematic annotation ^9–12^ have led to a current set of 466 mature *D. melanogaster* miRNAs ^13^.

Similar to other model organisms ^14^, only few fly miRNA deletions exert lethal phenotypes or strong morphological abnormalities. However, many miRNA have been found have subtle effects ^15,16^, which become more pronounced when the organism is challenged. It remains difficult to describe direct organismic miRNA effects via individual targets in a quantitative manner. Human tissue culture models have greatly enhanced our understanding about miRNA function, while our understanding of fly miRNA function is lagging behind. In *Drosophila*, a recent comparative study of small RNAs across 25 cell lines suggested that the miRNA landscape in non-ovary cell lines showed little diversity and low complexity in terms of relative expression levels of individual miRNAs, which would argue against fly cell lines as a good model to study miRNA function ^12^.

Although knowledge of mature miRNA sequences alone have enabled to identify physiologically relevant targets ^5,17^, computational methods have greatly contributed to successful miRNA target prediction, especially after recognition of the miRNA seed region (nt 2-7) ^18–21^. To date, there is a plethora of computational miRNA target predictions tools, including popular approaches such as TargetScan, MIRZA and mirSVR ^2,22,23^, which leverage conservation, target sequence context feature information or RNA::RNA hybridization energies and other features to improve prediction accuracy. Purely computational tools predict miRNA targets sites across entire 3’UTRs, neglecting cell type specific miRNA expression level and target site availability, which can lead to numerous and tightly spaced predictions. *In vivo* AGO binding information generated from Crosslinking and Immunoprecipitation (CLIP) followed by sequencing (CLIP-seq or HITSCLIP) methods has been used to greatly decrease the search space from whole 3’UTRs to about 30-40nt per AGO footprint ^24^. AGO footprints do not directly reveal the identity of the miRNA engaged, and in many cases, multiple possible miRNA seed matches overlap AGO binding sites. However, additional anchor points, such as the coverage summit ^24^ but especially the presence of diagnostic events (DE), rephrase the *in vivo* prediction problem to the assignment of the most plausible miRNA::mRNA pair within AGO footprints. DEs are introduced in the reverse transcription step during library generation and accumulate directly 5’ upstream of miRNA seed matches. PARCLIP enriches for abundant nucleotide conversions (i.e. T-to-C) in the sequenced read ^25^ but requires RNA-labeling with photoactivatable nucleosides (i.e. 4-Thiouridine). For HITSCLIP, nucleotide deletions have been mostly recognized to exhibit diagnostic potential ^26,27^. iCLIP on the other hand enriches for read truncations at the +1 nucleotide position of UV-crosslinked nucleotides ^28^. Dedicated computational methods leverage this biochemical, single nucleotide evidence of RBP::RNA interaction and have improved miRNA target identification to accuracy beyond solely sequence-based computational methods ^29,30^. Beyond assigning miRNA seed matches in relation to single nucleotide identifiers, chimeric miRNA::mRNA reads overlapping AGO footprints can be used to unambiguously identify the interacting miRNA ^31–34^.

Here we describe the absolute quantification of miRNAs in *Drosophila* S2 cells and find that miRNA expression landscapes in *Drosophila* cell lines are more complex than previously reported, owing to recent technological progress in small RNA cloning. We applied both HITSCLIP and PARCLIP to endogenous AGO1 protein, improving critical steps in the library cloning procedure, and we compared the predictive potential of single nucleotide diagnostic events (DEs) to assign ‘true’ miRNA::mRNA interactions. Making use of these features, we provide the first comprehensive transcriptome-wide map of miRNA target sites in fly. Using quantitative RNA-seq and sub-codon resolution ribosomal footprinting data in response to AGO1-depletion, we further functionally evaluated and validated different types of seed matches, confirming canonical miRNA functions. We suggest that fly cell lines are suitable models to study miRNA function and provide a fully quantitative resource with comprehensive transcriptome-wide miRNA binding sites and functional readouts.

## Results

### miRNA expression complexity in Drosophila S2 cells is greater than previously reported

In order to understand whether the low miRNA diversity previously observed in *Drosophila* cells ^12^ is indeed due to a low complexity in cell-type specific miRNA expression or in part due to miRNA detection limitations at that time, we generated new small RNA libraries for *Drosophila* S2 cells using adapters with randomized ends. Fixed adapter sequences had been identified as one of the major sources of miRNA quantification biases in small RNA sequencing experiments ^35,36^. Comparing mature miRNA sequences from both public and in-house S2 cell small RNA sequencing libraries (smRNA-seq) we found that miRNA expression values were more evenly distributed in samples generated using randomized adapter ends (Fig. 1A and Fig. S1A). While bantam-3p alone made up ∼60% of normalized miRNA reads in public smRNA-seq samples, it accounted for about ∼25% miRNA reads in our new samples. Other miRNAs, such as miR-14-3p and miR-7-5p, were detected at higher frequencies. These discrepancies are likely a result of miRNA detection differences between small RNA library cloning kits rather than differences in primary miRNA expression, as normalized RNA-seq coverage was unchanged between public and in-house RNA-seq libraries (Fig. 1B). Accordingly, we found that the read sequence composition at 5’ and 3’ read ends in public samples was noticeably skewed, possibly as a consequence of non-randomized adapter ends and concomitant pronounced ligation biases (Supplemental Fig. S1B). A noticeable proportion of small RNA reads from modENCODE-as well as newly generated samples aligned to common *Drosophila* viral genomes. Those reads likely represent 21nt long virus-derived siRNA and are unlikely to interfere with AGO1-mediated miRNA function (Supplemental note S1).

**Figure 1.**
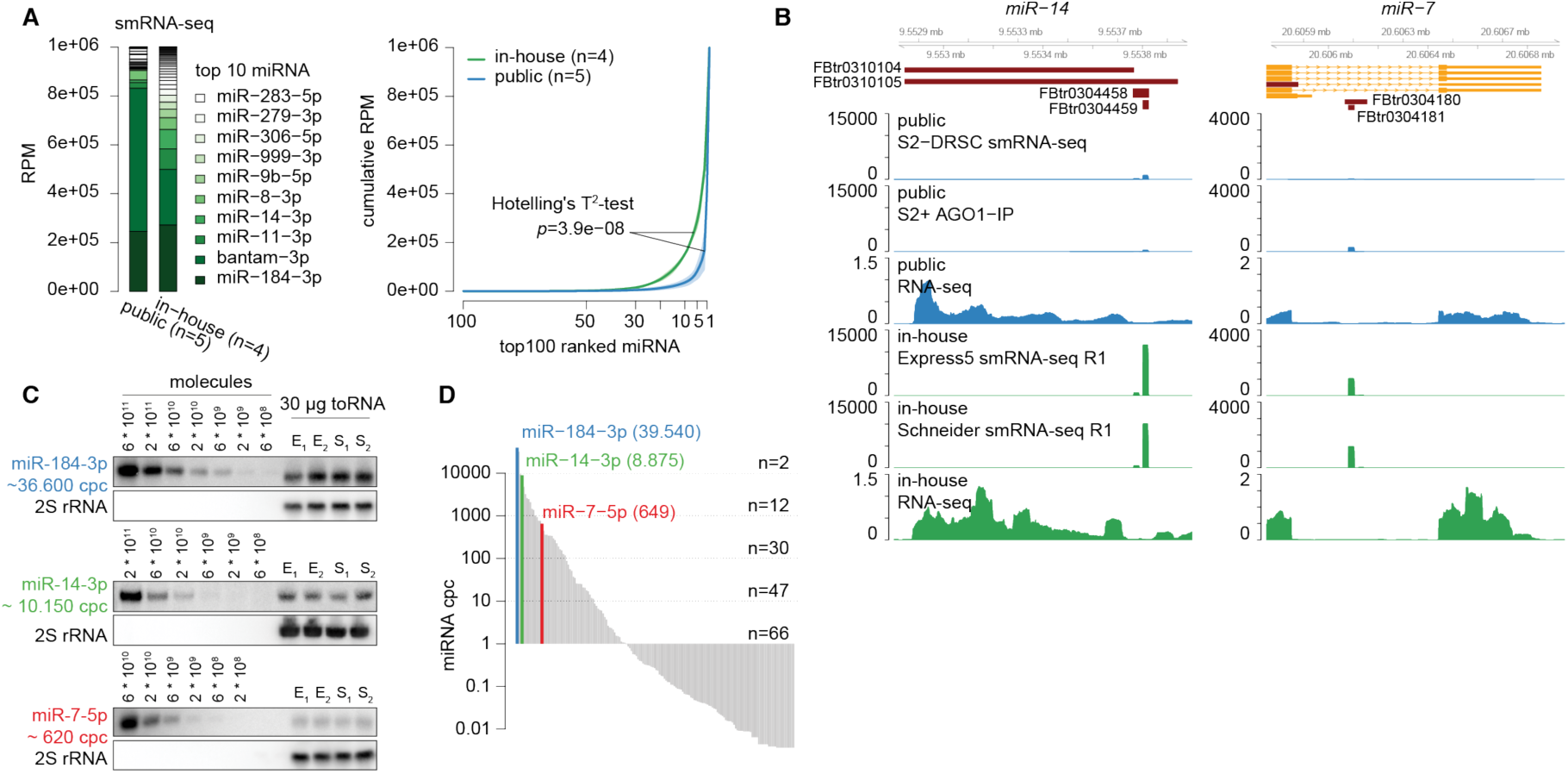
miRNA expression in Drosophila S2 cells is more complex than previously reported. A) miRNA quantification in publicly available and in-house smRNA-seq samples. miRNA annotated reads were normalized to reads per million (RPM). (Left) Barplot representing the mean RPM across replicates and sorted by in-house RPM. (Right) Cumulative miRNA RPM distribution of top 100 detected and RMP-ranked miRNAs. The solid line represents the mean across libraries, shades represent the standard deviation. B) Genome browser shot showing miR-14 and miR-7 reads and their respective RNAseq coverage at miRNA loci of representative libraries normalized to total library size. C) Quantitative miRNA northern blot for miR-184-3p, miR-14-3p and miR-7-5p, including their experimentally determined cpc. 2S rRNA served as a loading control for total RNA samples. D) Ranked distribution of fitted cpc values. Y-axis is in log10-scale.

Quantitative northern blot experiments confirmed that the previously lowly detected miR-14-3p was robustly detectable in S2 cells (Fig. 1C). We calculated miRNA copies per cell (cpc) for three miRNAs (miR-184-3p ∼36,600 cpc; miR-14-3p ∼10,150 cpc; miR-7-5p ∼620 cpc) and estimated cpc for all detected miRNA in smRNA-seq samples (Fig. 1D, supplemental table S1). Two miRNAs (miR-184-3p and bantam-3p) were present in more than 10,000 cpc and ∼30 miRNAs at more than 100 cpc. Taken together, miRNA expression levels in *Drosophila* S2 cells are more diverse than previously reported as a consequence of detection limitations.

### AGO1 HITS- and PARCLIP footprints harbor a similar set of single nucleotide DEs

To identify targets of the detected miRNAs, we performed two HITSCLIP ^37^ and two PARCLIP ^25^ experiments for endogenous AGO1 in S2 cells (Supplemental Fig. S2A and S2B). We updated individual library preparation steps and performed both CLIP-methods under similar conditions to be able to compare both approaches. Importantly, we replaced the RNase-T1 digestion with RNase-I digestion, which has no reported nucleotide cleavage bias, and again used 5’ and 3’ adapters with randomized ends to improve adapter ligation and help to efficiently remove PCR duplicates and, in part, sequencing errors. We sequenced all AGO1-CLIP amplicons close to estimated saturation resulting in 15,337,489 uniquely mapping reads (Supplemental Fig. S2C - S2E). Compared to human AGO2 PARCLIP libraries, we observed higher relative 3’UTR read density in fly cells (Supplemental Fig. S2F). This difference may be owed to a combination of higher density of predicted miRNA target sites in fly 3’UTRs compared to human 3’UTRs ^38^ and possibly differences in sequencing coverage. As instructive example, we confirmed all five originally predicted plus two additional bantam binding sites in the *hid* 3’UTR with AGO binding information from HITSCLIP and PARCLIP samples (Supplemental Fig. S2G) ^5^. Although harboring in total 45 predicted conserved and non-conserved 7/8mer seed matches for all detected miRNAs, only the predicted bantam seed matches were supported by the CLIP data.

The combination of AGO binding information and miRNA expression levels was highly effective to pinpoint the small set of actively engaged miRNA target sites from a large compendium of computationally determined candidates (Fig. 2A and 2B). We analyzed all CLIP data (both HITS- and PAR-CLIP) in the same framework for more comparability (see methods). First, we examined whether both CLIP methods would identify a similar set of AGO1 binding sites. Irreproducible discovery rate analysis indicated that both HITSCLIP and both PARCLIP replicates were characterized by high peak reproducibility, while reproducibility between both CLIP methods was less pronounced (Supplemental Fig. S2H). We pooled both HITSCLIP and both PARCLIP replicates and selected the *n* top peaks as indicated by an IDR < 0.25 (HITSCLIP n=8,971; PARCLIP n=11,667, supplemental tables S2 and S3) (Supplemental Fig. S2H). For both CLIP methods, 3’UTR annotating peaks were enriched relative to the number of peaks expected by chance (Supplemental Fig. S2I). IDR-selected AGO1 binding site positions were uniformly distributed within 3’UTRs, which is different from miRNA seed matches in human and in line with previous findings ^39^ (Supplemental Fig. S2J).

**Figure 2.**
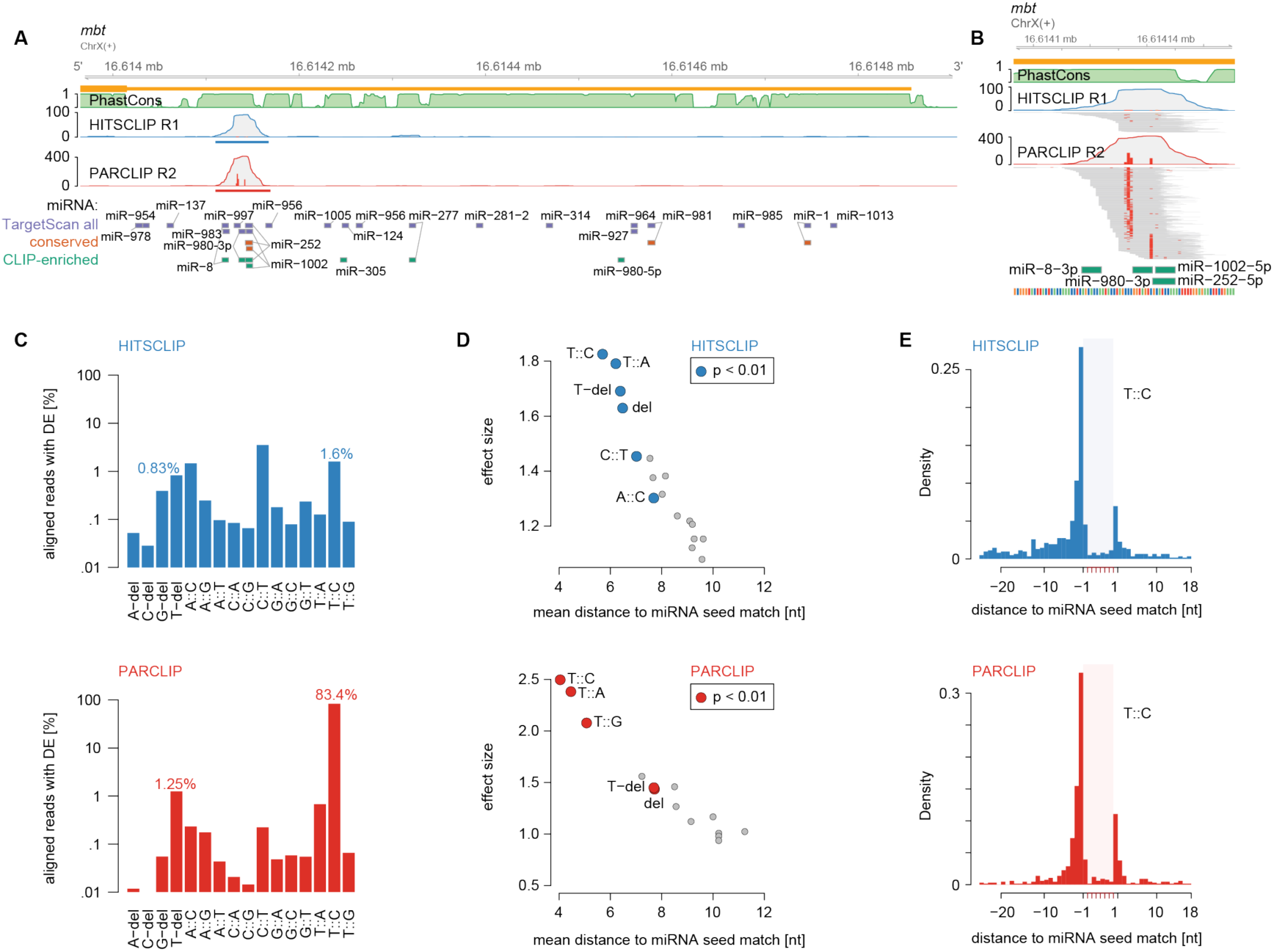
AGO1 HITSCLIP and PARCLIP diagnostic event comparison. A) Genome browser shot of the Drosophila gene mbt, depicting AGO1 HITSCLIP (blue) and PARCLIP (red) coverage tracks along its 3’UTR as well as 27way PhastCons scores (green). Blue and red bars indicate IDR-selected peak calls. Below, 7mer and 8mer seed matches for all miRNA in TargetScan 6.2 (conserved and non-conserved families), conserved miRNA (predicted conserved targets) and top59 CLIP-enriched miRNA (see supplemental Fig. S1A) are indicated. (y-axis shows the number of detected CLIP reads). B) Similar to A, genome browser shot of HITSCLIP and PARCLIP peak in mbt 3’UTR including alignments. Red squares in individual read alignments indicate T-to-C mismatches to the dm6 reference. Red bars within coverage tracks indicates the T-to-C conversion proportion at nucleotide resolution. Below, 7mer/8mer seed matches of CLIP-enriched miRNAs are indicated. C) Percentages of diagnostic events relative to all uniquely aligning reads. D) Results according to supplemental Fig. S2L. Scatterplot of mean distance to miRNA start (x-axis) relative to its effect size (y-axis). E) T-to-C conversion example according to D). Density of T-to-C conversion positional maxima relative to unique 7mer or 8mer matches in top 3000 IDR-selected 3’UTR peaks.

AGO footprints do not directly reveal which miRNA was bound. Several reports have exploited single nucleotide diagnostic events (DE) introduced during library preparation as additional anchor points within the AGO binding site, which give higher resolution information about direct RBP::RNA contacts. For AGO PARCLIP, T-to-C conversions have been found to be diagnostic to infer miRNA seed matches 3’ downstream ^25^. For AGO HITSCLIP, nucleotide deletions were the most recognized DE relative to miRNA seed matches ^26,27^. We found that PARCLIP peaks showed strong positional enrichment of T-to-C conversions, which is also observed in HITSCLIP peaks but to a lesser extent (Fig. 2B cf. ^40^.

We used the randomized adapter ends to filter aligned sequencing reads with mismatches to the reference genome to distinguish DE introduced at crosslinked nucleotides during reverse transcription from sequencing errors. After filtering, T-to-C conversions accumulated towards the middle of mapped reads for both PARCLIP and HITSCLIP samples (Supplemental Fig. S2K), in contrast to previous reports for mouse AGO2 CLIP ^26^. In AGO1 PARCLIP, more than 80% filtered uniquely aligning reads harbored T-to-C conversions (Fig. 2C). In AGO1 HITSCLIP data, we detected more reads with T-to-C conversions (1.6%) than reads harboring T-deletions (0.83%).

In order to evaluate the diagnostic potential of all possible nucleotide conversions and deletions, we evaluated the top 3000 3’UTR peaks in detail (Fig. 2D and S2L; see methods). For both, PARCLIP and HITSCLIP T-to-C conversions preferentially peaked 5’ proximal to unique 7- and 8-mer seed matches within AGO1 footprints (Fig. 2E). Although PARCLIP T-to-C conversions were by far more abundant, the less frequent conversions in HITSCLIP can nevertheless indicate crosslinked nucleotide 5’ proximal to seed matches. In PARCLIP, not only T-to-C conversion, but also T-to-A, T-to-G conversions and T-deletions occur closer to seed matches than expected by chance (Fig. 2D and S2M). In HITSCLIP, T-to-C, T-to-A conversions and T-deletions showed similar preference. (Fig. 2D and S2M). About 80% of the top 3000 AGO1 HITSCLIP 3’UTR peaks contained at least one T-to-C conversion, while T-deletions occurred in less than 25% and showed slightly less diagnostic potential. For both AGO1 PARCLIP and HITSCLIP, crosslinked nucleotides are best indicated by T-to-V conversions and T-deletions, though at different frequencies.

### T-centric DEs enable efficient miRNA target site prediction in PAR- and HITSCLIP

To assess the impact of T-to-V conversions together with T-deletions, we used microMUMMIE ^29^, a hidden Markov model that integrates CLIP binding profiles and their DEs with sequence matches to predict miRNA seed matches within AGO1 binding sites. For both CLIP methods we chose peaks with at least 2 DEs (hereafter referred to as cluster ^41^). In AGO1 PARCLIP almost all IDR-selected 3’UTR peaks contain at least 2 T-to-C conversions (n=3740/3890 3’utr peaks). In AGO1 HITSCLIP more than 50% (n=1661/3086 3’utr peaks) of the IDR-selected 3’UTR peaks were clusters based on T-to-V or T-del DE, while T-to-C conversions accounted for more clusters than T-deletions (Fig. 3A).

**Figure 3.**
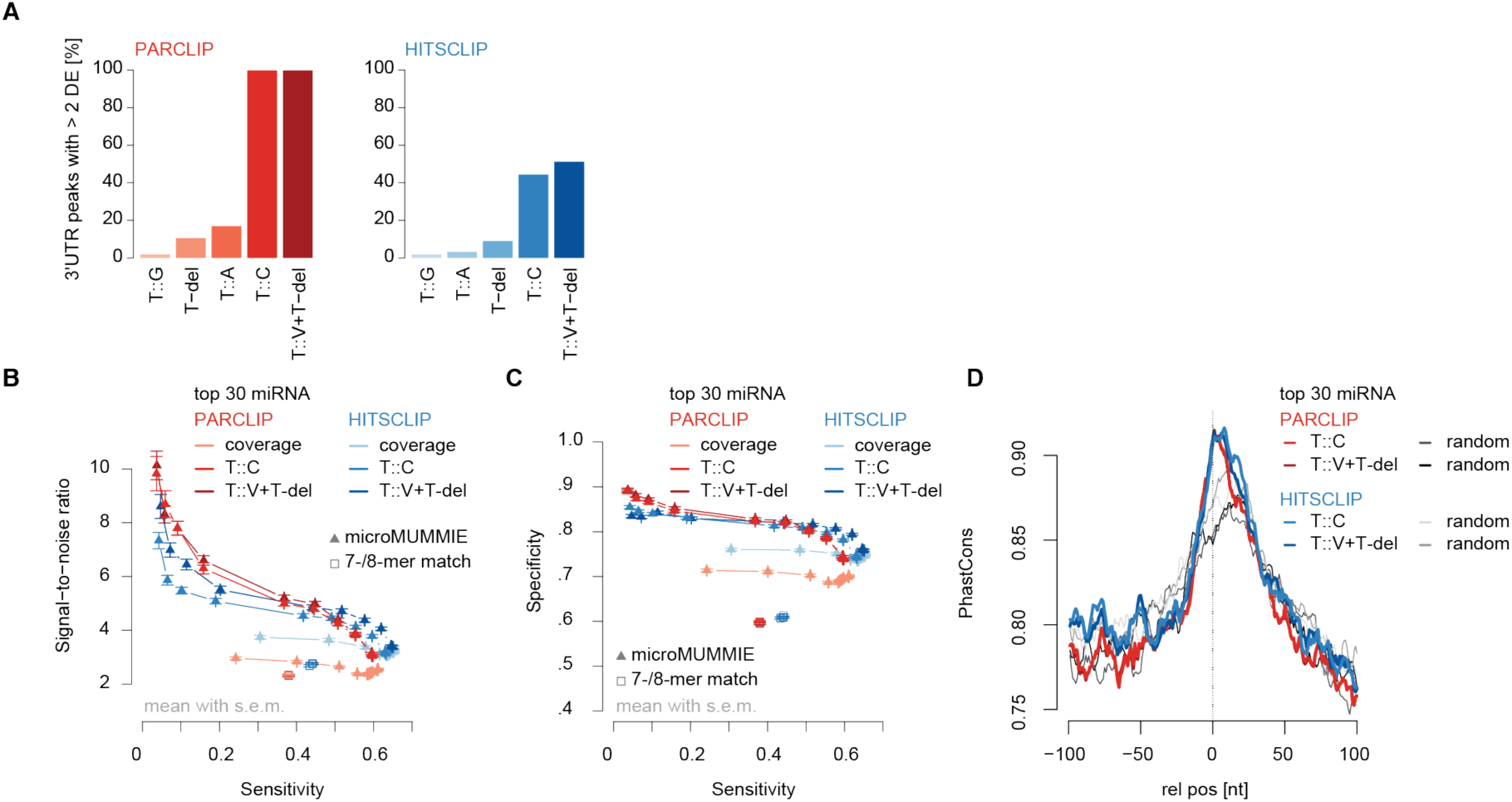
microMUMMIE assigned miRNA seed matches on PARCLIP and HITSCLIP. A) Proportion of IDR-selected peaks forming clusters (≥ 2 DE per 3’UTR peak) depending on individual or combined DEs. B) SNR estimate for HISTCLIP and PARCLIP derived DE signal for miRNA seed match predictions given the top 30 CLIP-enriched miRNA relative to 30 shuffled decoy miRNA. In each case, the top 1500 clusters were used. The results are depicted as mean across 100 individual shuffling experiments, with error bars representing SEM. Individual triangles indicate changes in chosen microMUMMIE variance levels. Squares show basic 7mer-A1, 7mer-m8 or 8mer-A1 matches anywhere within clusters. X-axis depicts sensitivity. Coverage = inferred single nucleotide peak summit position. C) Similar to B) but depicting specificity vs. sensitivity. D) UCSC 27way PhastCons scores relative to the inferred crosslinked nucleotides for Clusters with miRNA seed match (at microMUMMIE variance 0.01; Viterbi mode) prediction or a random nucleotide within the same peak.

We ran microMUMMIE on the top 1500 PARCLIP clusters harboring T-to-C conversions and predicted miRNA seed matches for the miRNAs that were detected in CLIP samples relative to same number of decoy miRNAs (see methods). CLIP samples showed a clear bimodal miRNA read distribution, suggesting that the top 59 miRNAs are actively engaged in AGO1-RISC complexes (referred to as comprehensive miRNA set) (Supplemental Fig. S1A). We found that a smaller set of top 30 detected miRNAs had the best trade-off maintaining high signal-to-noise ratio (SNR), while maintaining almost maximal sensitivity (referred to as high-confidence miRNA set) (Supplemental Fig. S3A and S3B). Comparing the predictive potential of DEs between both CLIP methods using the high-confidence miRNA set, miRNA::mRNA pairs were assigned with higher SNR at lower sensitivity values in AGO1 PARCLIP-derived clusters as compared to HITSCLIP-derived clusters (Fig. 3B and 3C; Fig S3C and S3D). HITSCLIP clusters may thus harbor a higher proportion of true positive miRNA seed matches compared to PARCLIP, but the high density of PARCLIP-derived DEs has a higher predictive value. In all cases, using DEs within the top 3’UTR clusters were more predictive of real miRNA seed matches than using the position of the peak summit (coverage midpoint) (Fig. 3B/3C and Fig. S3C/S3D). While combining T-to-V or T-del DEs helped in the case of HITSCLIP, PARCLIP clusters did show similar SNR and sensitivity using T-to-C conversions only. For both methods, we found a similarly strong increase of PhastCons conservation scores relative the inferred crosslinked (Fig. 3D and S3E). In summary, we predicted miRNA seed matches for AGO1 PARCLIP and HITSCLIP clusters at comparable SNRs. However, DEs were detectable as a function of sequencing depth, and their prevalence is much lower especially in AGO1 HITSCLIP peaks with lower coverage.

### Canonical miRNA binding sites function via 3’UTRs targeting

To confirm miRNA function, we knocked down AGO1 expression using double-stranded RNA (dsRNA) mediated gene silencing and performed mRNA sequencing and ribosomal footprinting relative to control treatments (Supplemental Fig. S4A and Fig. S4B). We calculated mRNA expression changes, changes in ribosomal footprinting, and translational efficiency (Supplemental table S4). We assessed whether identified AGO1 binding sites in different regions of mRNAs had similar effects. For IDR-selected peaks in both CLIP-methods we found that repression alleviation upon AGO1-depletion was strongest for genes bound in 3’UTRs (Supplemental Fig. S4C). Changes in RNA levels and ribosomal footprinting data were concordant for the majority of AGO1-bound targets (Supplemental Fig. S4D). It has been suggested previously that AGO-binding-dependent translational repression precedes RNA degradation ^42^ and that AGO-binding in coding regions may specifically influence target gene translational efficiency ^43^. In our data, only a small subset of AGO1 targets bound in their 3’UTR were characterized by additional changes in translational efficiency that were not explained by mRNA abundance changes (Supplemental Fig. S4C). We also did not observe strong changes in translational efficiency for genes targeted in coding regions relative to genes without AGO1 binding sites. However, our data was derived from 72h dsRNA knock-down and thus may not be well suited to address preceding changes in translational efficiency.

As expression changes were most pronounced for 3’UTR bound AGO1 targets, we focused on providing a reference miRNA target site annotation to binding sites in this annotation category. Since miRNA seed match prediction on PARCLIP T-to-C conversions had the best SNR, and DE prevalence was much higher than in AGO1 HITSCLIP, we reanalyzed the PARCLIP data using the PARCLIP-tailored peak caller PARalyzer ^41^ (Supplemental tables S5-S7). First, in order to explain as many AGO1 3’UTR clusters as possible, we pooled both PARCLIP samples and predicted miRNA seed matches for the 59 CLIP-enriched miRNAs (referred to as comprehensive miRNA target site map; Supplemental table S8). Similar to previous studies, not all AGO1 binding sites can be explained by a canonical miRNA seed match. In addition to spurious non-functional interactions in genomic cross-linking data sets, this fraction may consist at least partially of AGO1-binding sites without canonical miRNA seed match that may still be able to function (bulge sites, 3’ compensatory sites, center sites, etc. ^44–46^. However, the prevalence of such sites is still largely unclear, and we found that target gene expression of genes with clusters lacking canonical seed matches was not noticeably different from non-targeted genes (Supplemental Fig. S4E). Furthermore, slight changes could be also explained by canonical seed matches of miRNAs not included in the comprehensive miRNA set, as well as targeting in other transcript regions.

For lower ranked clusters of the pooled AGO1 PARCLIP data sets, prediction certainty was gradually reduced (Supplemental Fig. S4F). In order to arrive at a high-confidence miRNA target site map, we predicted miRNA seed matches for the top 30 CLIP-enriched miRNA that showed good sensitivity, while maintaining high SNR on the IDR-selected peaks (referred to as high-confidence miRNA target site map; Supplemental table S10). Here, we predicted miRNA seed matches on both AGO1 PARCLIP samples separately and kept reproducible miRNA seed match predictions. The gold standard comprises 5026 miRNA seed match predictions on 2601 expressed genes (Fig. 4A). These reproducible predictions showed stronger target repression alleviation upon AGO1 knockdown than genes with non-reproducible target sites as part of the comprehensive miRNA target site set (Supplemental Fig. S4G).

**Figure 4.**
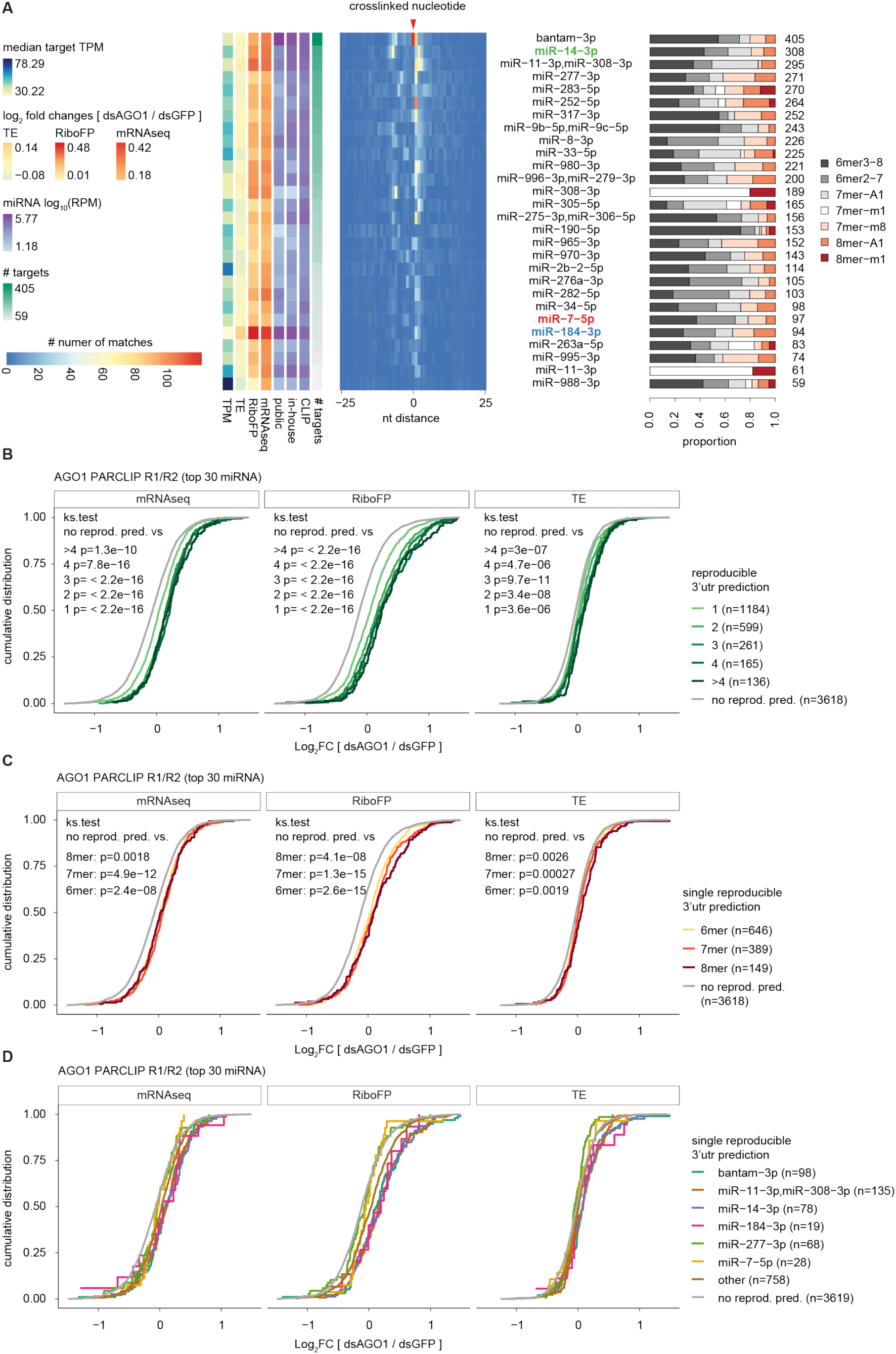
Functional evaluation of canonical miRNA seed match predictions. A) Heatmap showing positional miRNA prediction prevalence relative to the identified crosslinked nucleotide for the top 30 CLIP-enriched miRNAs within PARalyzer-derived 3’UTR clusters. Only miRNA seed match prediction reproducible in both AGO1 PARCLIP replicates were considered. miRNAs are ranked by the number of predicted targets. The proportion of seed match types is shown on the right. On the left, the medians of steady state target expression levels (TPM), log2 fold changes of dsAGO1 vs. dsGFP treated samples for TE, RiboFP and mRNA-seq are shown for all miRNA targeted genes, followed by the mean miRNA RPM expression levels in public and in-house smRNA-seq as well as CLIP data sets. Results shown were derived at microMUMMIE variance 0.01 using viterbi mode. B) Cumulative distribution of mRNA-seq, RiboFP and TE log2 fold changes for genes with 1, 2, 3, 4 or more than 4 reproducible miRNA seed match predictions relative to genes without reproducible predictions. P value was calculated in a two-sided Kolmogorov-Smirnov test versus genes without reproducible miRNA seed match predictions. C) Similar as in B) but isolating genes with exactly one reproducible miRNA seed match prediction stratified by 6mer, 7mer or 8mer binding mode. D) Similar as in C) but depicting log2 fold changes for individual miRNAs (miR-184-3p, miR-14-3p and miR-7-5p compared to three other miRNAs with the most miRNA predictions).

### S2 cell miRNA repertoire manifests terminally differentiated state

The number of predicted targets correlated well with CLIP-derived miRNA quantification (Pearson correlation coefficient 0.47, p=0.012). Accordingly, we found (the previously lowly detected and most CLIP-enriched miRNA) miR-14-3p to have the second-most reproducible miRNA target sites. On the other hand, miR-184-3p was associated with comparably few targets and did not follow this general relationship. Yet, those few targets exhibited strong repression alleviation upon AGO1 knockdown. The number of reproducible miRNA predictions had a strong cumulative effect (Fig. 4B), which was stronger than differences in miRNA seed match types (Fig. 4C, S4H). For some miRNAs, miRNA effects indeed increased in the order of 6mer < 7mer < 8mer (i.e. bantam-3p, miR-184-3p), but we also found examples of abundant miRNAs showing the opposite relationship (i.e. miR-277-3p). Individual miRNAs therefore differed from each other in target suppression strength and mode (Fig. 4D), possibly a sign of miRNA:mRNA target stoichiometry differences.

Having information about *in vivo* bound miRNA target sites in S2 cells provides the unique opportunity to describe the collective miRNA targetome and individual miRNA modules. We found 1237 being targeted by a combination of at least two miRNAs, while 1364 genes harbor one single miRNA binding site (Fig 5A, inset). We noted that all unique miRNA target sets are larger than any miRNA pair, suggesting that no larger specific combinatorial target gene sets exist (Fig 5A). To test whether these rather unique target sets address distinct biological processes, we checked for the presence of specific gene function (GO) categories. We could identify a group of strongly enriched GO terms around fly development, morphogenesis, signaling and cell-to-cell communication (Fig. 5B; Supplemental file S2), processes suggested to be prime miRNA targets in *Drosophila* ^47^. This group showed a very similar enrichment pattern across most individual miRNA target sets, suggesting a high overlap in their targeted developmental processes. Indeed, calculating semantic GO-term similarities supports the notion that the majority of miRNA targets share similar GO-term enrichments (Fig. 5C).

**Figure 5.**
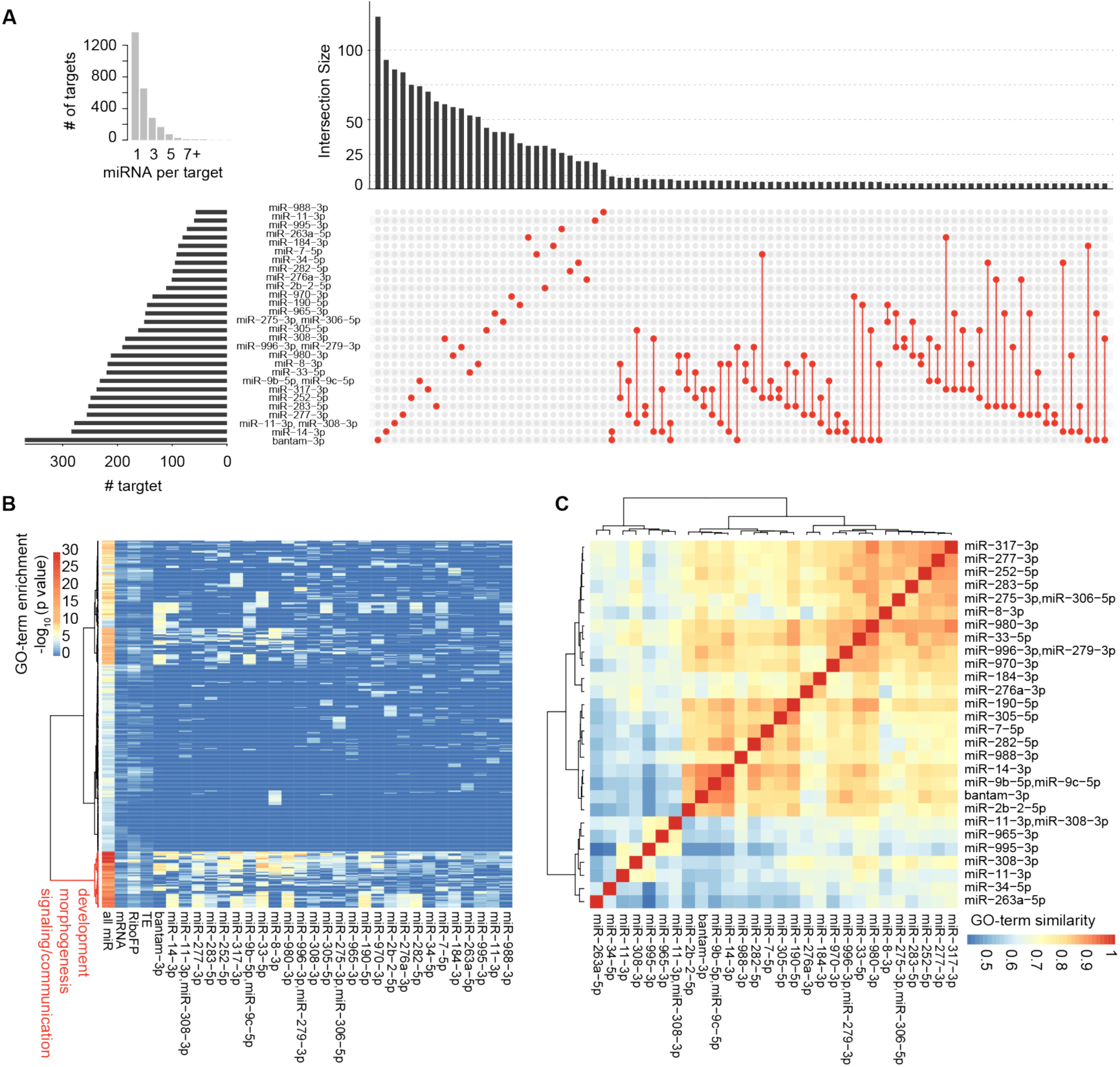
miRNAs in S2 cells collectively target genes involved in development. A) Overview of S2 miRNA targetome. The inset on the left shows the number of detected genes with unique (=1) to up to 13 reproducible 3’UTR miRNA binding sites. Upset plot showing all possible miRNA target overlaps with minimally 4 shared genes (n= 88 combination >= 4 genes). Bar plot on the left indicates the number of all targets per miRNA. The bar plot on top indicates the size of the unique target set. The largest target gene sets exist for individual miRNAs. The largest intersect for co-targeting miRNA has a size of 9 targets. The sets are indicated by red dots, connected by red lines. B) Biological process gene ontology (GOBP) enrichment for all miRNA targets (all miR: n=2,601), top decile of genes upregulated on mRNA level upon AGO1-depletion (mRNA), top decile of genes upregulated on ribosomal footprinting level upon AGO1-depletion (RiboFP), top decile of genes upregulated Translational efficiency level upon AGO1-depletion (TE; each n=597), and all individual miRNA target sets, relative to all genes considered during functional analysis previously (n=5,962). All significantly enriched (p < 0.001; fisher’s exact test; n=501) GO-terms for all miRNA targets were selected, merged to the corresponding enrichments in all other sets, and row-wise clustered (distance = maximum, clustering function = ward) after p value −log10-transformation, resulting in two main clusters. miRNAs are sorted by the number of targets. We did not observe enriched GO terms for individual miRNA target sets, which were not already covered by enrichments in all miR. (Supplemental file S2). C) Pair-wise GO-term similarities using GOSemSim (Yu et al. 2010), for the top 100 enriched GOBP terms given p < 0.001 (fisher’s exact test), and clustered (distance = euclidean, clustering = ward).

## Conclusion

Despite its importance as a model system, the fly community has been lacking a comprehensive, quantitative, in vivo map of *D. melanogaster* miRNA targets. To fill this gap, we describe a resource of cellular miRNA copy numbers, comprehensive miRNA target sites, as well as functional response data including ERCC spike-in RNA-seq and matched sub-codon resolution ribosomal footprinting data utilizing randomized adapters. Here, we focused our efforts on comparing the miRNA target prediction potential for AGO1 CLIP methods, and the evaluation of miRNA function. Together, we support that at least Schneider S2 cells lines can serve as a valuable model to study fly miRNA function.

We found the expressed miRNA pool to be more diverse than reported. T4 RNA ligases, most commonly used during small RNA cloning, were shown to have sequence biases and/or nucleic acid secondary structure hindrance in ligating single stranded RNA or DNA oligos, which can lead to miRNA mis-quantification of multiple orders of magnitude ^35,48–50^. Randomizing adapter ends for miRNA cloning can efficiently reduce those biases, and results showed good agreement between complementary miRNA quantification methods ^36^. Beyond overcoming ligation limitations, randomized adapter ends serve furthermore as unique molecular identifiers (UMIs), which help to distinguish individual ligation events from duplications introduced during PCR or sequencing. This is especially critical for low-complexity smRNA-seq libraries, where often thousands of identical reads align to only a few miRNA loci. Accordingly, we show that this new miRNA expression data is in better agreement with quantitative Northern blots than previous libraries prepared without randomized adapter ends. This finding may not be limited to Drosophila S2 cells and plausibly extends to all of the 25 fly cell lines recently profiled for modENCODE ^12^.

For some miRNAs, detected expression levels changed drastically between public and new smRNA-seq quantification. MiR-14-3p has early on been associated with an anti-apoptotic phenotype in fly ^6^ and since then been implicated in multiple other regulatory cues ^15,51–54^. Early smRNA cloning and pyrophosphate sequencing as well as SOLiD-sequencing already indicated mir-14 to be abundant in S2 cells ^9,55^, but in all but one of the public smRNA libraries analyzed mir-14 was significantly underrepresented as compared to our new smRNA-seq data (Supplemental Fig. S1A). We found miR-14 levels as one of the most engaged miRNA in AGO1-RNP complexes, targeting the second most genes following bantam.

While we, and others ^25^, observed a general correlation of miRNA expression level and the number of predicted miRNA targets (Fig. 4A), miR-184 did not follow this trend. We mapped fewer target sites than expected from its expression, and its target genes were on average more strongly derepressed upon AGO depletion and showed a relatively strong effect on translational regulation (Fig. 4A and 4D). MiR-184 has been previously found to be required for embryonic axis formation and has an age dependent effect on female germline development ^15,56^. It has also been found responsive to high-sucrose treatment in fly and mouse as well as in diabetic mouse models ^57,58^, suggesting a conserved response mechanism. In both cases, the miR-184 levels are reported to drop quickly and strongly upon treatment and disease state, while miRNA expression changes are known to be normally modest. The quick drop suggests a short miRNA half-life. Given a high miRNA:mRNA target ratio in S2 cells, effective target regulation would require strong changes in miRNA levels. It seems therefore tempting to speculate that miR-184 shows common strong regulation as a result of high miR-184::target mRNA ratios in fly and mouse.

AGO binding information greatly enhances accuracy in assigning the miRNA:mRNA gene pairs ^24,25^, as the search space for short miRNA seed matches is reduced dramatically from whole 3’UTRs to AGO footprints. If ambiguity remains, single nucleotide diagnostic events can be used for assigning the right miRNA ^26–30^. DEs are known for all three major CLIP protocols (PARCLIP, HITSCLIP, iCLIP) but it has so far been unclear how the diagnostic potential of CLIP-type specific DEs compare. In agreement with previous reports, we found T-nucleotide DEs (conversions and deletions) for miRNA seed matches located 3’-downstream for both HITSCLIP and PARCLIP ^40^, but these DEs (especially T-to-C conversions) were much more abundant in PARCLIP. Contrary to previous reports ^26^, HITSCLIP T-to-C conversions showed higher diagnostic potential than T-deletions, due to multiple possible reasons: A) The original study used less stringent mapping parameters (∼75% of reads contain conversions), possibly shadowing a lower fraction of informative diagnostic events. B) Mapping of short reads including mismatches remains specifically challenging and differs across aligners ^59^, and the ∼19x smaller fly genome (dm6 vs. mm10) implies higher mapping confidence. C) Our use of randomized UMI adapter ends enabled us to identify and remove sequencing errors from aligned reads. A similar approach has been used for AGO iCLIP samples ^34^. Overall, we found the combination of HITSCLIP T-to-V conversions and T-deletions to lead to competitive SNR, sensitivity and specificity to predict miRNA seed matches, but only for the top 1500 peaks. Given that only 2.5 % (1 in 40 reads) of uniquely aligned reads contain such DE, only peaks with substantial coverage can be used for this analysis, while this limitation does not exist in PARCLIP. For HITSCLIP peaks without DEs, the coverage mid-point could still act as anchor point ^24^, but with lower SNR ^29^. Importantly, our observations can be leveraged for *in vivo* endogenous AGO1 CLIP experiments, where the 4SU incorporation of 4SU into transcripts may be difficult or impossible.

We increased the count of experimentally supported miRNA target sites in *D. melanogaster* from currently 12 (Diana TarBase v7.0 ^60^) and 150 (miRTarBase 7.0 ^61^), respectively, to more than 5000 reproducible sites. It is possible that miRNA targeting follows slightly different rules in different clades. For example, in *C. elegans* additional miRNA targeting modes were found to be more conserved than expected by chance (6mer-A1 and 8mer-1U) ^38^. Moreover, miRNA seed matches in fly do not occur preferentially towards 3’UTR start and end ^39,62^, which is supported by our AGO1 binding data and possibly a consequence of drastically shorter 3’UTR length in flies ^38^. As 3’UTRs can undergo extensive lengthening for example in the fly nervous system and thus increase cis-regulatory 3’UTR space ^63,64^, this picture may depend on the tissue or for individual miRNAs. Furthermore, local AU-content was found to be predictive for miRNA target sites in human ^2^, but the 3’UTR AU-content is higher in fly and thus may be less predictive. Beyond facilitating a quantitative model of miRNA targeting in fly, our comprehensive target map is thus an excellent starting point to improve Drosophila target prediction.

## Methods

### miRNA quantification

For miRNA quantification, we considered AGO CLIP and smallRNA-seq alignments after the step of multimapper removal. This was chosen for two main reasons: 1) miRNA harbor large proportions of untemplated 3’-end modification resulting in mis-matches towards read-ends that do not result from sequencing or adapter trimming errors. 2) Public smallRNA-seq libraries (Supplemental file S1) were generated without introducing UMIs, which are required for UMI-based sequencing error removal.

Reads annotating to mature miRNA were quantified and normalized to reads per million (RPM) by dividing by the total number of miRNA-annotating reads and multiplication with 1*10^6^. CLIP-enriched miRNAs were identified fitting a two-component mixture model ^65^ to the log_10_-transformed and RPM-normalized miRNA counts as a mean across CLIP libraries.

To infer copies per cell (cpc) for all miRNA detected, we first fit a linear regression model to the experimentally determined cpc and in-house smallRNA-seq derived mean RPM after log-transformation. The resulting model was used to predict cpc for all detected miRNA.

Experimentally determined miRNA cpc values fitted better with miRNA reads per million (RPM) derived from in-house smRNA-seq libraries (R-squared = 0.999, p=0.0013, residual std. er. = 52.4), than with public data sets (R-squared = 0.95, p=0.14, residual std. er. = 5810). Accordingly, fitted cpc values for all miRNA were more coherent with in-house smRNA-seq derived RPM (in-house - R-squared = 0.989, residual std. er. = 3030; public - R-squared = 0.932, residual std. er. = 11600).

### HITSCLIP and PARCLIP of endogenous AGO1 protein

AGO1 HITSCLIP and PARCLIP experiments were performed in biological replicates, originally described in ^25^ with the following changes. Buffers were used from ^66^. For AGO1 PARCLIP, culturing medium was supplemented with 400 µM 4-Thiouridine (4-SU) (*SIGMA #T4509*) 17 hours overnight before harvest. 4-SU incorporation was determined to be approximately half as efficient as in HEK293T cells determined by thiol-specific biotinylation dot-blot assays as described previously ^67^. Semi-adherent cells were scraped and washed in ice-cold PBS prior to 254 nm or 365 nm UV-irradiation (400 mJ/cm^2^), respectively. Liquid cell pellets were snap frozen in liquid nitrogen and stored at −80°C until further usage. For library preparation, cells were thawed on ice and quickly lysed in NP40 lysis buffer (50 mM Tris–HCl pH 7.4, 100 mM NaCl, 1 % Igepal CA-630 (NP40), 0.1 % SDS, 0.5 % sodium deoxycholate, Complete Protease Inhibitor to a final concentration of 2x, RNAsin 40 U/ml lysis buffer; 1 ml lysis buffer per approximately 0.3*10^9^ cells, 1.2*10^9^ to 1.3*10^9^ cells in total per sample). After treatment with RNaseI (1:333 v/v or 300 U/ml lysate for HITSCLIP and 1:400 v/v or 250 U/ml lysate for PARCLIP samples, because of slight concentration differences between lysate; TurboDNase (4 U/ml) for 3 min at 37 °C and 1100 rpm, immunoprecipitation (IP) was carried out with polyclonal AGO1 (*Abcam #ab5070*, 20 µg per sample) coated magnetic protein A dynabeads (*Life Technologies #10002D*)(100 µl) on spin-cleared cell extracts for 2 hrs at 4 °C.

After IP, CLIP samples were stringently washed three times with high salt buffer (50 mM Tris–HCl, pH 7.4, 0.666 M NaCl, 1mM EDTA, 1% Igepal CA-630 (NP40), 0.1% SDS, 0.5% sodium deoxycholate), followed by PNK-buffer washes. Samples were radioactively 5’end-labelled with γ-^32^P-ATP including a subsequent addition of 1 µl high-molar ATP to the reaction for efficient 5’end phosphorylation. The crosslinked protein-RNA complexes were resolved on a 4-12 % Bis-Tris-Polyacrylamid gel. The SDS-PAGE gel was transferred to a nitrocellulose membrane and the protein-RNA complexes migrating at an expected molecular weight were excised. RNA was isolated by Proteinase K treatment and phenol-chloroform extraction, ligated to 3’ adapter and 5’ adapter (Supplemental file S1), reverse transcribed using Superscript III *(Life Technologies #18080044)*, PCR-amplified (PCR cycles: HITSCLIP 20 cycles; PARCLIP 19 cycles), and gel-purified. Note, after the 3’Adpater ligation step, each AGO1 sample was split into approximately 19-24 nt (miRNA fraction) and 24-35nt (target fraction) long fragments, cloned, amplified and sequenced separately. The amplicons were sequenced single-end as a multiplexed pool on HiSeq2000 (Illumina) with 51 cycles.

### CLIP and smallRNA library processing

For AGO1 HITSCLIP and PARCLIP libraries, sequencing reads from 19-24 nt and 24-35 nt fraction were combined before processing. For all fly CLIP libraries we quality-filtered reads using the fastx-tool kit [-q 10 -p 95] (http://hannonlab.cshl.edu/fastx_toolkit/), and adapter-trimmed using cutadapt v1.8 ^68^ [--overlap=3; -m 24] (Supplemental file S1), discarding untrimmed reads. Reads were collapsed (duplicate removal) still including the 4 randomized nucleotides at both ends of the sequencing read. Randomized adapter ends got trimmed after read collapsing and added to the read identifier for further usage and treated as unique molecular identifiers (UMIs). As the smaller fly genome allowed higher mapping rates, we required minimally 16 nt read length. rRNA mapping reads were removed prior to aligning to the fly genome. We filtered multimapping reads and only kept the best alignment of a read if the second best alignment had more than one mismatch more than the best alignment. SmallRNA samples were processed accordingly. If no randomized adapter ends (UMIs) were present, we did not apply PCR duplicate removal. miRNA quantification on CLIP libraries was done after this processing step. Further, we filtered out all reads with mismatches relative to the genome in the first and last two nucleotides. Next, we removed reads with mismatches relative to the genome reference which were likely introduced during sequencing and thus represent sequencing errors and not diagnostic events. For this, we grouped alignments based on genomic coordinates (Chr, start, end, strand) and UMIs. In the case where alignments shared all coordinates and harbored the same UMI, while differing from each other and/or the reference sequence, we sorted by copy number (retained from read collapsing) and removed reads with relative lower copy number and higher mismatch prevalence to the local high copy number reference read.

For the comparative CLIP analysis we called peaks using Piranha v1.2.1 ^69^ [- s -b 20 -a 0.95 -v]. To work around Piranha’s assumption that the smaller genomic coordinate is the read start irrespective of the strand, we called peaks on the read midpoints. For spliced reads, the read midpoint was assigned to the part of the read with the more extensive exon overlap. Peak reproducibility was estimated using Irreproducible Discovery Rate (IDR) ^70^ on the peak read counts with an overlap ratio of 0.1. For AGO1 CLIP libraries, we chose an IDR < 0.25 for reproducing peaks between CLIP replicates and selected the top *n* peaks (HITSCLIP pooled n=8,971; PARCLIP pooled n=11,667).

AGO1 PARCLIP data was additionally processed using PARalyzer ^41^ embedded in the PARpipe wrapper pipeline (https://github.com/ohlerlab/PARpipe) as described before ^71^. In brief, pre-processing included the steps of adapter trimming, PCR duplicate removal as described above. Randomized adapter nucleotides were trimmed using Flexbar (Dodt et al., 2012). Here, reads were mapped using bowtie requiring minimally 20nt read length. Removal of rRNA reads, sequencing errors and multimapper were not applied here but left to the pipelines default setting. For group and cluster calling, PARalyzer v1.5 parameter setting were set to default except requiring minimally five unique reads to initiate a group call, while neglecting PCR-duplicate information. PARalyzer-generated clusters were filtered for T-to-C conversion specificity of at least 0.6 and higher.

### Determination of diagnostic event positional preferences

The top 3000 3’UTR annotated IDR-selected Piranha peaks were selected for both, AGO1 HITSCLIP and PARCLIP and extended on either side by 5 nt. Within peak sequences, we searched for miRNA seed matches (7mer-A1, 7mer-m8 or 8mer-A1) for the 20 most abundant miRNA in CLIP and 1000 times the same number of dinucleotide-shuffled miRNA using the TargetScan.pl script v6.1 ^72^. Shuffled decoy miRNAs were generated using uShuffle ^73^ on mature miRNA sequence and rejecting decoy sequences if they overlapped an non-shuffled (referred to as true) miRNA seed within the top 20 miRNAs. We selected peaks with exactly one seed match. Individual diagnostic event (DE) tracks at single nucleotide resolution (i.e. all T-to-C conversion) were isolated from the CLIP alignment files and mapped relative to true miRNA or decoy miRNA seed matches in a window of ±25 nt from the genomic miRNA seed match start. For each window around a miRNA seed match the position with maximal DE occurrence was determined.

For each DE, we calculated the mean distance of maximal occurrence to the seed match start across all windows for true miRNAs and shuffled miRNAs sets. Similarly, we calculated the ratio of 1/Gini-coefficient to determine positional enrichments with variable distance to miRNA seed match starts. Empirical significance was assigned with p < 0.01, if less than one percent of the 1000 individual shuffle experiments yielded lower mean distance or higher 1/Gini values than the true miRNAs. The effect size was calculated forming the ratio of the sample median of all mean distances generated by shuffling experiments and mean distance for true miRNAs.

### microMUMMIE signal-to-noise, sensitivity and specificity estimation

miRNA target prediction evaluation was conducted in three scenarios: 1) To assess the optimal number of miRNA to query. 2) To compare microRNA target prediction between AGO1 HITSCLIP and PARCLIP. 3) To evaluate miRNA target prediction with respect to the relative rank of miRNA clusters. In each experiment, microMUMMIE was used without the option of including TargetScan-provided branch length scores shown to improve prediction accuracy in human ^29^. Branch length score cut-offs for a dm6-based multiple sequence alignment have not been determined in a same way yet. Available branch length score cut-offs for a dm3-based 12way multiple sequence alignments, did not improve prediction SNR.

To evaluate miRNA target prediction between AGO1 HITSCLIP and PARCLIP we isolated DE for T-to-C conversions or the combination of T-to-A, T-to-C and T-to-G conversions as well as T-deletions (referred to as T-to-V+T-del) from fully filtered alignment files. Peaks with at least two diagnostic events were considered as clusters. Cluster boundaries were refined by trimming its edges if coverage dropped below 5 reads. As described in the PARalyzer method ^41^ we applied kernel density smoothing to the DEs within each cluster. Similarly, we determined the coverage summit. Like this, large parts of PARalyzer, including its output formats (distribution files storing smoothed DE information) were implemented in R relying Bioconductor packages ^74^. For the top 1500 3’UTR clusters (width/read count) in AGO1 HITSCLIP and PARCLIP, we estimated the miRNA seed match prediction accuracy using microMUMMIE, as described in the microMUMMIE methods ^29^. In brief, we ran microMUMMIE using the top *n* CLIP-enriched miRNA plus the same number of dinucleotide-shuffled decoy miRNA. Shuffling was done using uShuffle ^73^. Decoy miRNAs were rejected, if their seed nucleotides 3-7 were overlapping with any true miRNA nucleotide 3-7 sequence to avoid 6mer overlaps. Only miRNA seed match predictions overlapping input clusters were retained. Predictions overlapping several transcript isoforms or miRNA seed family members were collapsed to single genomic coordinates. The signal-to-noise ratio (SNR) describes the number of true miRNA seed match predictions divided by the number of decoy miRNA seed match predictions. Sensitivity is defined as number of clusters with at least on true miRNA seed match prediction, while specificity is defined as the ratio of true miRNA seed match predictions divided by the number of all (true and decoy miRNA) predictions. We ran microMUMMIE in viterbi mode and without conservation at 10 variance levels (var = 1.5, 1, 0.75, 0.5, 0.25, 0.2, 0.15, 0.1, 0.01, 0.005), depicting the mean SNR, sensitivity and specificity with its standard error of the mean (SEM) for 100 individual shuffling and training experiments. Branch lengths scores calculated for the dm6-centric 27-way multiple sequence alignment provided by UCSC did not improve microMUMMIE predictions, most likely as suitable branch length score cut-offs were not available.

Similarly, we estimated miRNA target prediction for AGO1 PARCLIP libraries processed with PARalyzer. Clusters were ranked as described above and binned into groups of 1000 clusters, before calculating SNR, sensitivity and specificity for each bin separately.

For a conservative and comprehensive set of miRNA target site predictions microMUMMIE was run on PARalyzer-derived 3’UTR clusters from both AGO1 PARCLIP libraries separately and only reproducible predictions were retained. MicroMUMMIE was run at 6 different stringency levels (variance var = 0.5, 0.25, 0.2, 0.15, 0.1 and 0.01).

Further detailed method descriptions can be found in supplemental methods.

## Supplemental data contains tables for

table S1 miRNA quantification

table S2 AGO1 HITSCLIP IDR-selected peaks

table S3 AGO1 PARCLIP IDR-selected peaks

table S4 Xtail results [dsRNA AGO1-low / dsRNA GFP]

table S5 PARalyzer-Cluster on AGO1 PARCLIP R1 sample

table S6 PARalyzer-Cluster on AGO1 PARCLIP R2 sample

table S7 PARalyzer-Cluster on pooled AGO1 PARCLIP samples

table S8 Comprehensive miRNA target set (pooled PARCLIP; top59 miRNA; 3’utr)

table S9 MiRNA target set (reproducible PARCLIP R1/R2; top59 miRNA; 3’utr)

table S10 High-confidence miRNA target set (reproducible PARCLIP R1/R2; top30

table S11 MiRNA target set (reproducible PARCLIP R1/R2; top30 miRNA; cds)

table S12 MiRNA target set (reproducible PARCLIP R1/R2; top30 miRNA; 5’utr)

Supplemental file S1 summarizes RNA and DNA oligos, processed data sets and ERDN/SQRC smallRNA spike-in. Supplemental file S2 contains GO-term enrichments.

### Acknowledgements

We gratefully acknowledge the kind gift of ERDN/SRQC smallRNA spike-in reagents from the Timo Breit lab (University of Amsterdam). The anti-PABP antibody was kindly provided by Marina Chekulaeva (Max-Delbrück-Center for Molecular Medicine, Berlin). H.H.W. and U.O. were supported in part by National Institutes of Health (NIH) grant R01GM104962 and HFSP grant RGY0093/2012.

## Author contributions

H.H.W., N.M. and U.O. conceived the experiments and computational analysis. H.H.W., S.L. and A.H. executed the experiments. H.H.W. did the computational analysis. H.H.W. and U.O wrote the manuscript.

